# Domain-architecture aware phylogenetic profiling indicates a functional diversification of type IVa pili in the nosocomial pathogen *Acinetobacter baumannii*

**DOI:** 10.1101/2023.02.01.526601

**Authors:** Ruben Iruegas, Katharina Pfefferle, Stephan Goettig, Beate Averhoff, Ingo Ebersberger

## Abstract

The Gram-negative bacterial pathogen *Acinetobacter baumannii* is a major cause of hospital-acquired opportunistic infections. The increasing spread of pan-drug resistant strains, paired with a surge in virulence, makes *A. baumannii* top-ranking among the ESKAPE pathogens for which novel routes of treatment are urgently needed. Comparative genomics approaches have successfully identified genetic changes that coincide with the emergence of pathogenicity in *Acinetobacter*. Genes that are prevalent both in pathogenic and a-pathogenic Acinetobacter species were not considered ignoring that virulence factors may emerge by the modification of evolutionarily old and widespread proteins.

Here, we increased the resolution of comparative genomics analyses to also include lineage-specific changes in protein domain architectures. Using type IVa pili (T4aP) as an example, we show that two pilus components, FimU and the pilus tip adhesin ComC, exist with markedly differing domain architectures in pathogenic *Acinetobacter* species. ComC, has gained a von Willebrand Factor type A domain displaying a finger-like protrusion, and we provide experimental evidence that this finger conveys virulence-related functions in *A. baumannii*. Both genes together form an evolutionary cassette, which has been replaced at least twice during *A. baumannii* diversification. The resulting strain-specific differences in T4aP layout suggests differences in the way how individual strains interact with their host. Our study underpins the hypothesis that *A. baumannii* uses T4aP for host infection as it was shown previously for other pathogens. It also indicates that many more functional complexes may exist whose precise function can be rapidly adjusted by changing the domain architecture of individual proteins.

**Author Summary:** Type IVa pili (T4aP) are hair-like, extendable, and retractable appendages that many bacteria use for interacting with their environment. Several human pathogens have independently recruited these pili for processes related to host infection, but the modifications necessary to turn T4aP into virulence factors are largely unknown. Here, we studied if and how T4aP components have changed in the nosocomial pathogen *A. baumannii* compared to its largely a-pathogenic relatives in the *Acinetobacter* genus. Most *A. baumannii* isolates have T4aP with a pilus tip adhesin containing a protein domain not seen outside the pathogenic clade. This domain is essential for bacterial motility and contributes to host cell adhesion and natural competence. However, some isolates have T4aP resembling those of largely a-pathogenic species in this genus. This indicates that the way these pili are used during infection processes differs between *A. baumannii* isolates probably as a result of niche adaptation. In a broader perspective, our findings highlight that many relevant genetic differences between pathogens and their a-pathogenic relatives emerge only on the domain- and sub-domain level. This suggests that comparative genomics studies have only uncovered the tip of the iceberg of genetic determinants that contribute to *A. baumannii* virulence.

## Introduction

*Acinetobacter baumannii* is a Gram negative nosocomial pathogen that accounts for about 5% of the total bacterial infections worldwide [1–3]. The world-wide spread of multi- or even pan-resistant *A. baumannii* isolates [4–6] is accompanied by a surge in virulence [6–10], and thus novel therapeutic treatments are necessary for a sustainable infection management. Over the past years, experimental studies integrated with comparative genomics analyses have therefore sought to identify genetic determinants of *A. baumannii* virulence [11–23]. The identified factors are involved into a broad spectrum of biological processes including NOS and ROS resistance and metabolic adaptation, but some also indicate changes in the way how the pathogen interacts with its environment [17, 20].

Pili, or ‘hair-like’ surface appendages, are main mediators of bacterium-environment interaction [24]. Most bacterial phyla possess type IV pili (T4P) [25, 26], multi-purpose nanomachines that act via dynamic cycles of extension and retraction mediated by cytoplasmic motor ATPase-driven polymerization and depolymerization of pilin subunits [27–32]. To date, three sub-types of T4P are known of which the sub-type ‘a’ is most prevalent [26]. T4aP are involved in a variety of functions [24], of which surface adhesion, bacterial motility and the uptake of environmental DNA are tightly connected to bacterial virulence [33, 34]. It is thus not surprising that several Gram negative and Gram positive human pathogens use T4aP for processes connected to host infection [33, 35–40]. In *Acinetobacter*, T4aP play a role in cell adhesion [41], twitching motility and natural transformation [30, 42–44]. Moreover, the two-component regulatory system BfmRS, which is important for survival of *A. baumannii* in a murine pneumonia model, also controls T4aP production [45]. While this suggests that T4aP could also drive *A. baumannii* virulence, the most comprehensive comparative genomics study so far between pathogenic and a-pathogenic *Acinetobacter* species failed to detect differences that could hint towards lineage-specific changes in T4aP formation or function [20].

T4aP are prevalent in bacteria irrespective of their lifestyle [24]. Their recurrent recruitment by pathogens for processes connected to host infection therefore suggests that only considerably subtle modifications are necessary to transform T4aP into a virulence factor. Therefore, any adaptive changes in pathogenic *Acinetobacter* might have escaped the attention thus far, because they require higher resolving analyses beyond determining the presence/absence of T4aP components. Indeed, structural variants of the major pilin subunit PilA were recently detected in *A. baumannii* strains. This suggested the existence of functionally diverse T4aP in this species [41, 46], but a comprehensive analysis on this level of resolution for all T4aP components considering, at the same time, the wealth of sequence data covering the full range of *Acinetobacter* diversity is missing. Therefore, it is still unclear to what extent T4aP differ between members of this genus and what consequences this may have for the interaction of *A. baumannii* with the human host.

Here, we set out to chart the extent of variation in the precise design of T4aP across the genus *Acinetobacter.* We integrated genus-wide phylogenetic profiles of T4aP components across more than 800 bacterial isolates with an assessment of domain architecture dissimilarity as a proxy of functional divergence. This revealed that the pseudopilin FimU and the pilus tip adhesin ComC exist in two different domain architecture layouts and together form an evolutionary cassette that was exchanged at least twice during *A. baumannii* diversification. The ComC variant that is prevalent in *A. baumannii* and in closely related pathogens of the Acinetobacter calcoaceticus/baumannii complex (ACB complex) differs from its orthologs in other *Acinetobacter* species by the presence of a N-terminal von Willebrand-factor type A Pfam domain (VWA_2). 3D structure modelling revealed the presence of a finger-like protrusion that likely originated from a lineage-specific insertion into the structural scaffold of the von Willebrand-factor type A domain. We showed experimentally that the VWA_2 domain is essential for adhesion to HUVECs and for twitching motility, and at least contributes to natural transformation. Deleting only the finger-like protrusion had the same effect as the deletion of the entire VWA_2 domain indicating that it is essential for the function of this ComC variant. Taken together, our results provide for the first-time evidence that pathogenic *Acinetobacter* have modulated the precise function of T4aP by changing the structural layout of the pilus tip adhesin.

## Results

### Phylogenetic profiles of Type IV pilus components

*Acinetobacter* T4aP components are best characterized in the naturally transformable bacterium *A. baylyi* ADP1 (Fig. 1A; [42]). We included the prepilin peptidase PilD because the corresponding gene is part of the *pilBCD* operon in *A. baylyi,* and because it is likely involved in T4aP biogenesis [43, 47]. All *A. baylyi* ADP1 T4aP components are represented in the *A. baumannii* type strain *Ab* ATCC 19606^T^ (Table 1), which allowed to reconstruct the evolutionary history of T4aP from an *A. baumannii* perspective. A targeted ortholog search for the individual *Ab* ATCC 19606 ^T^ T4aP components across 855 *Acinetobacter* isolates covering the diversity of the genus complemented with 29 outgroup species resulted in the phylogenetic profiles shown in Fig 1B. Orthologs to all T4aP components are present throughout the genus *Acinetobacter*, and most are also found in the outgroup species. Using ComC as an example, we showed that a replacement of the stringent ortholog search with a more inclusive, but less specific unidirectional BlastP search identified the pilus tip adhesin PilC1 as the unique best hit in *N. meningitidis* (S1 Fig). This indicates that a limited sensitivity of the ortholog search explains at least part of the gaps in the phylogenetic profiles of T4aP components [48].

**Figure 1.**
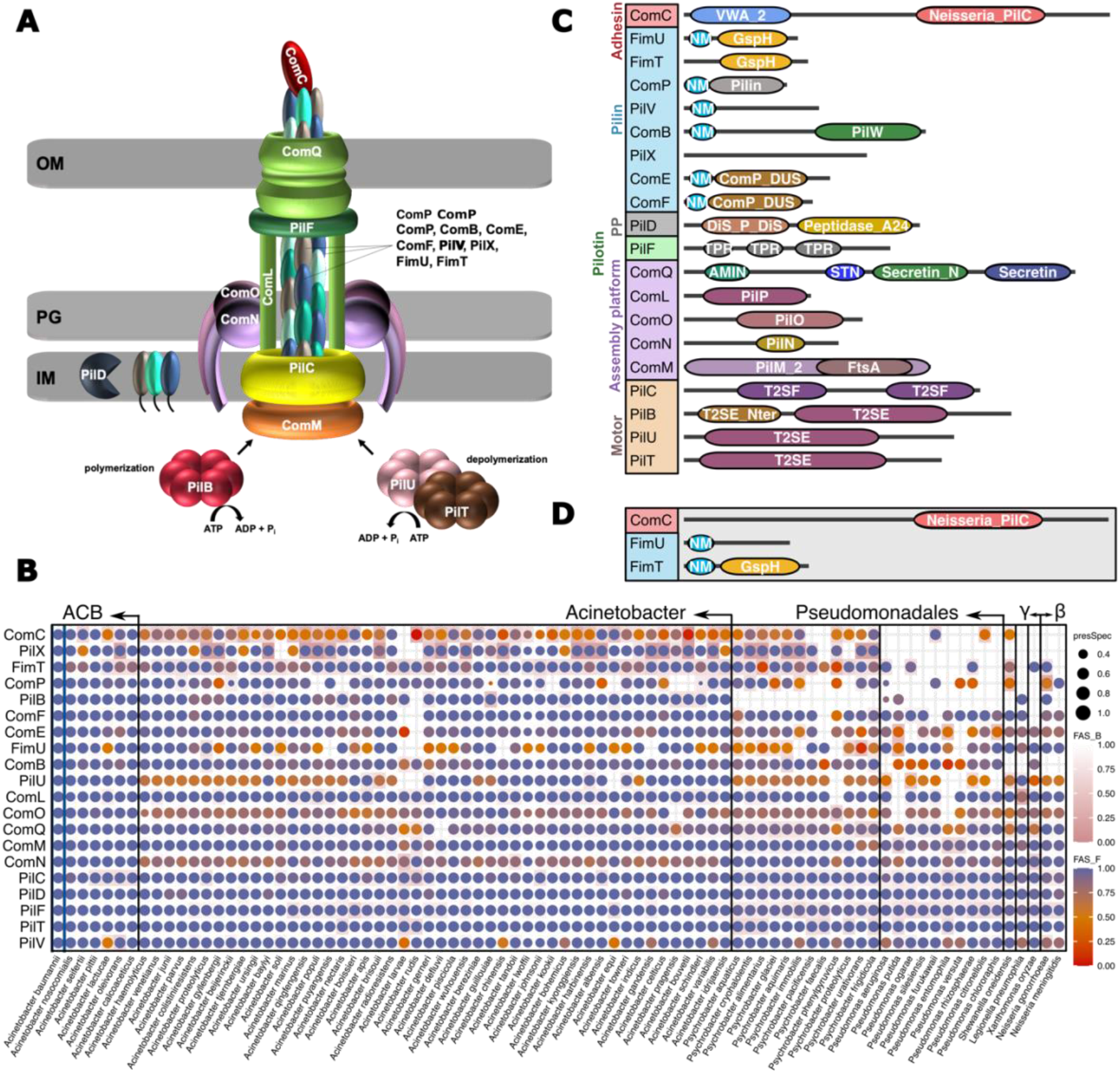
Characterization of *Acinetobacter* type IVa pilus components. (A) Model of the type IVa pilus (T4aP) in *A. baylyi*. OM – Outer membrane; IM – Inner membrane; PG – peptidoglycan. The individual components are listed in Table 1. (B) Phylogenetic profiles of the *Ab* ATCC 19606 ^T^ T4aP components across 884 bacterial isolates representing 83 species. Taxa are summarized on the species level. A dot indicates the presence of an ortholog in the respective species, and the dot size represents the fraction of the subsumed isolates an ortholog was identified in. The color encodes the median domain architecture similarity (FAS score) between the protein in *Ab* ATCC 19606 ^T^ and its orthologs using the *Ab* ATCC 19606 ^T^ protein as reference (FAS_F; dot color) or the ortholog (FAS_R; cell color). The color gradients from blue to orange (dot color), and from white to pink (cell color) indicate decreasing domain architecture similarities. Taxa are ordered according to increasing phylogenetic distance to *A. baumannii.* ‘γ’ and ‘β’ represent *γ*- and *β proteobacteria,* respectively. The full data is available as S1 data. (C) Pfam domain architectures of the T4aP components in *Ab* ATCC 19606 ^T^. Protein lengths are not drawn to scale. Pfam accessions and domain descriptions are available in S1 Table. PP – Prepilin peptidase; NM – N_Methyl. (D) Most common alternative domain architectures for ComC, FimU and FimT represented in (C). Representative architectures are shown for *A. baumannii 1297* (ComC and FimU) and for *A. baylyi* (FimT).

**Table 1.** Genes involved into T4aP formation in *A. baylyi* ADP1 and in *A. baumannii* ATCC 19606 ^T^.

|  | *A. baylyi*<br>ADP1 | *A. baumannii*<br>ATCC 19606<sup>T</sup> | Locus<br>start | Locus<br>end | Annotation |
| --- | --- | --- | --- | --- | --- |
| *pilU* | WP_004922051.1 | WP_000347036.1 | 124856 | 125974 | PilT/PilU family type 4a pilus ATPase |
| *pilT* | WP_011182164.1 | WP_000220756.1 | 126002 | 127069 | type IV pilus twitching motility protein (pilus ATPase) |
| *pilF* | WP_011182074.1 | WP_000537691.1 | 636841 | 637608 | type IV pilus biogenesis/stability protein PilW |
| *pilB* | WP_004920473.1 | WP_001274985.1 | 831958 | 833670 | type IV-A pilus assembly ATPase PilB |
| *pilC* | WP_004920476.1 | WP_000279216.1 | 833700 | 834926 | type II secretion system F family protein |
| *pilD* | WP_004920478.1 | WP_001152283.1 | 834926 | 835786 | prepilin peptidase |
| *comM* | WP_004923715.1 | WP_000537766.1 | 1511699 | 1512721 | pilus assembly protein PilM |
| *comN* | WP_004923716.1 | WP_000201227.1 | 1512721 | 1513362 | PilN domain-containing protein |
| *comO* | WP_004923719.1 | WP_000076101.1 | 1513359 | 1514099 | type 4a pilus biogenesis protein PilO |
| *comL* | WP_004923726.1 | WP_000695065.1 | 1514110 | 1514637 | pilus assembly protein PilP |
| *comQ* | WP_004923730.1 | WP_001017037.1 | 1514700 | 1516865 | type IV pilus secretin family protein |
| *comP*<sup>S</sup> | WP_004923779.1 | WP_000993713.1 | 1529663 | 1530091 | pilin |
| *fimU* | WP_004923829.1 | WP_001214059.1 | 1539389 | 1539862 | prepilin-type N-terminal cleavage/methylation domain-containing protein |
| *pilV* | WP_004923832.1 | WP_002194578.1 | 1539856 | 1540416 | type IV pilus modification protein PilV |
| *comB* | WP_004923834.1 | WP_000079195.1 | 1540417 | 1541418 | PilW family protein |
| *pilX* | WP_004923837.1 | WP_086221418.1 | 1541475 | 1542233 | pilus assembly protein |
| *comC* | WP_004923840.1 | WP_085940514.1 | 1542341 | 1546099 | VWA domain-containing protein |
| *comE* | WP_004923843.1 | WP_001046417.1 | 1546112 | 1546594 | prepilin-type N-terminal cleavage/methylation domain-containing protein |
| *comF* | WP_004923844.1 | WP_000788344.1 | 1546591 | 1547016 | prepilin-type N-terminal cleavage/methylation domain-containing protein |
| *fimT* | WP_004922618.1 | WP_000477156.1 | 1756167 | 1756682 | type II transport protein GspH |

\$ synonym: *pilA*

### Domain architecture changes in T4P components

The phylogenetic profiles provide no evidence for a lineage-specific modification of T4aP within *Acinetobacter* that is driven by the gain or loss of individual genes. We therefore increased the resolution by comparing the domain architectures of the individual *Ab* ATCC 19606 ^T^ T4aP components to those of their orthologs (Figs 1B-D). For most proteins, domain architectures are either conserved across the genus, or differ only in the presence/absence of low complexity regions or coiled coil regions (see S2 Fig). However, the architectures of four proteins, PilX, FimT, FimU, and ComC deviate to an extent between orthologs that it could indicate a change in function. Of these proteins, two are unlikely to drive T4aP diversification on a larger scale. PilX orthologs differ by the sporadic absence of a PilX Pfam domain (PF13681), which probably reflects chance effects affecting the accurate annotation of the Pfam domain (Fig S3). Domain architecture difference between FimT orthologs are caused by the presence of an N-terminal Methyl_N domain (PF07963) with only few exceptions, among them *Ab* ATCC 19606 ^T^ (Figs. 1D and S4 Fig A-B). The domain loss in the *A. baumannii* type strain is caused by a substitution that results in a tyrosine at a position that is typically occupied by a leucine or an isoleucine in PF07963 (S4 Fig C-D). While this change is very likely of functional relevance, it is specific to individual strains. More compelling are the findings for FimU and the pilus tip adhesin ComC. Both proteins exist in two main domain architecture variants in the genus (Fig 1D and S5 Fig). FimUVar1 differs from FimUVar2 by the presence of a GspH Pfam domain (PF12019). This domain is characteristic for pseudopilins which are involved in bacterial type II export systems. ComC_Var1_ differs from ComC_Var2_ by the presence of an N-terminal von Willebrand factor type A domain (Pfam VWA_2; PF13519).

### Evolutionary histories of ComC and FimU in *Acinetobacter*

We next investigated the evolutionary histories of ComC and FimU in greater detail. A phylogenetic analysis of the ComC orthologs revealed that the species *A. baumannii* and the genus *Acinetobacter* are both paraphyletic regarding this locus (Fig 2 and S6 Figure), and an alternative tree where the *Acinetobacter* isolates were forced into monophyly could be rejected (AU test; p=0.002; [49]). In the genome of *Ab* ATCC 19606 ^T^, *comC* and *fimU* reside in close vicinity (see Fig 3). To determine whether the two genes are not only physically but also genetically linked, we labeled each taxon in the ComC tree with the respective variant combination for FimU and ComC. Within *A. baumannii*, but also for most isolates from the *Acinetobacter calcoaceticus-baumannii* (ACB) complex, ComC_Var1_ is almost exclusively found together with FimUVar1, and ComC_Var2_ is typically associated with FimUVar2 (Fig 2 and S7 Fig). The association of the variants is only broken up in early branching *Acinetobacter* species outside the ACB complex. Interestingly the tree reveals a third clade comprising individual *A. baumannii* isolates and one representative from *A. calcoaceticus*. The domain architectures of FimU and ComC in this clade resemble that of Var1 on first sight, but differences emerge on the sub-domain level (see below). To give credit to its distinct phylogenetic placement, we refer to it as Var1-2 to distinguish it from the more abundant Var1-1.

**Figure 2.**
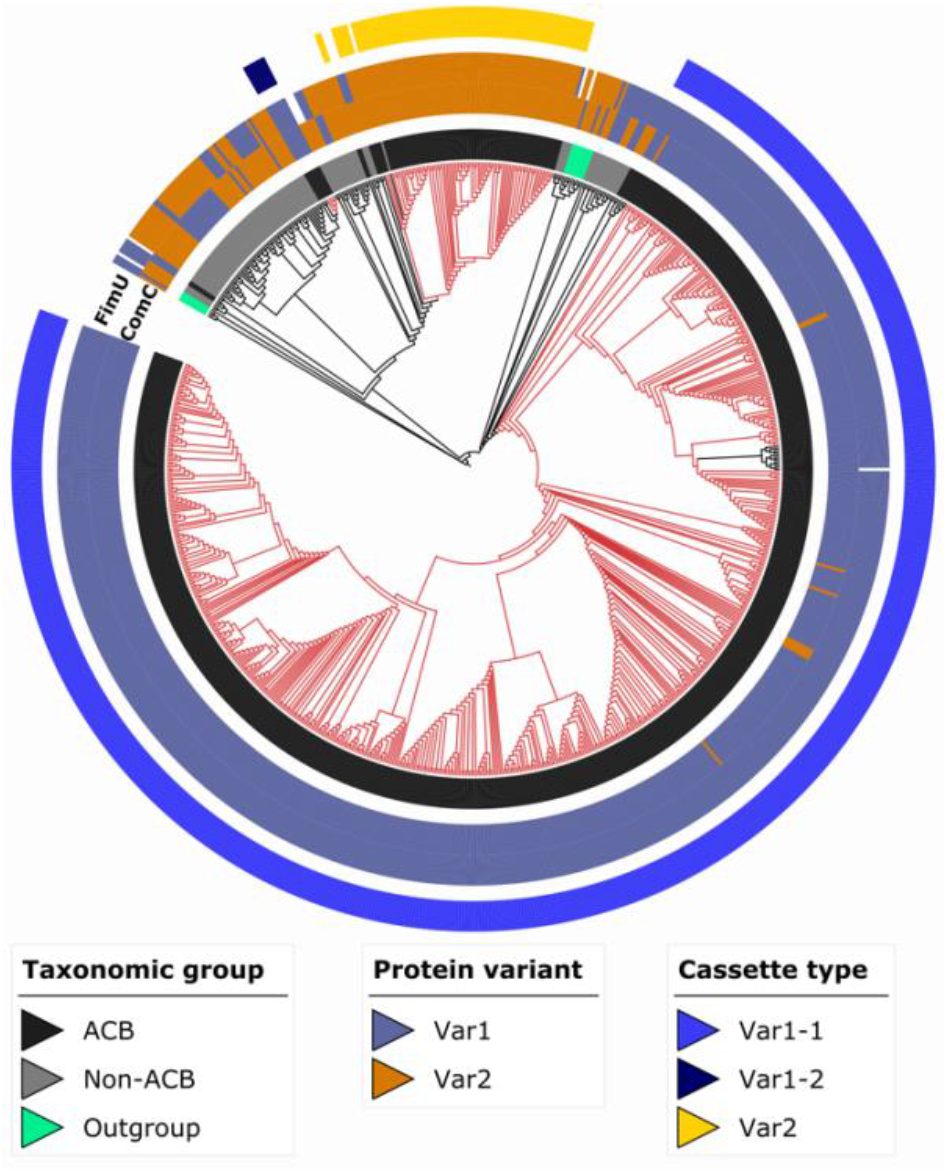
Maximum likelihood phylogeny of ComC. Red branches indicate *A. baumannii* isolates. The inner ring (ring 1) identifies the taxonomic group the isolates in the tree are assigned to. The variant assignments of FimU and ComC across the investigated taxa are provided in rings 2 and 3, respectively. The three evolutionary cassettes present in the ACB clade (see main text) are represented in the outermost ring 4 (see main text for further details). A high-resolution version of this tree is given in S6 Figure. ACB – *Acinetobacter calcoaceticus-baumannii* complex.

The phylogenetic analysis has revealed that ComC and FimU are represented by three evolutionarily distinct lineages with two alternative Pfam domain architecture layouts in *A. baumannii*. This indicates that recombination has affected the evolution of this locus, and we next assessed the length of the genomic region that was likely involved in these recombination events. We focused on a region covering 5 kbp upstream and 3.5 kbp downstream of ComC, which harbors five additional genes involved in T4aP formation as well as six flanking genes with different functions (Fig 3). To rule out that changes in gene order represent a physical barrier to recombination, we confirmed that the order of these 13 genes is conserved across the *Acinetobacter* diversity (S8 Fig). The analysis revealed that the recombination block spans all seven T4aP components in this region but excludes the flanking genes.

**Figure 3.**
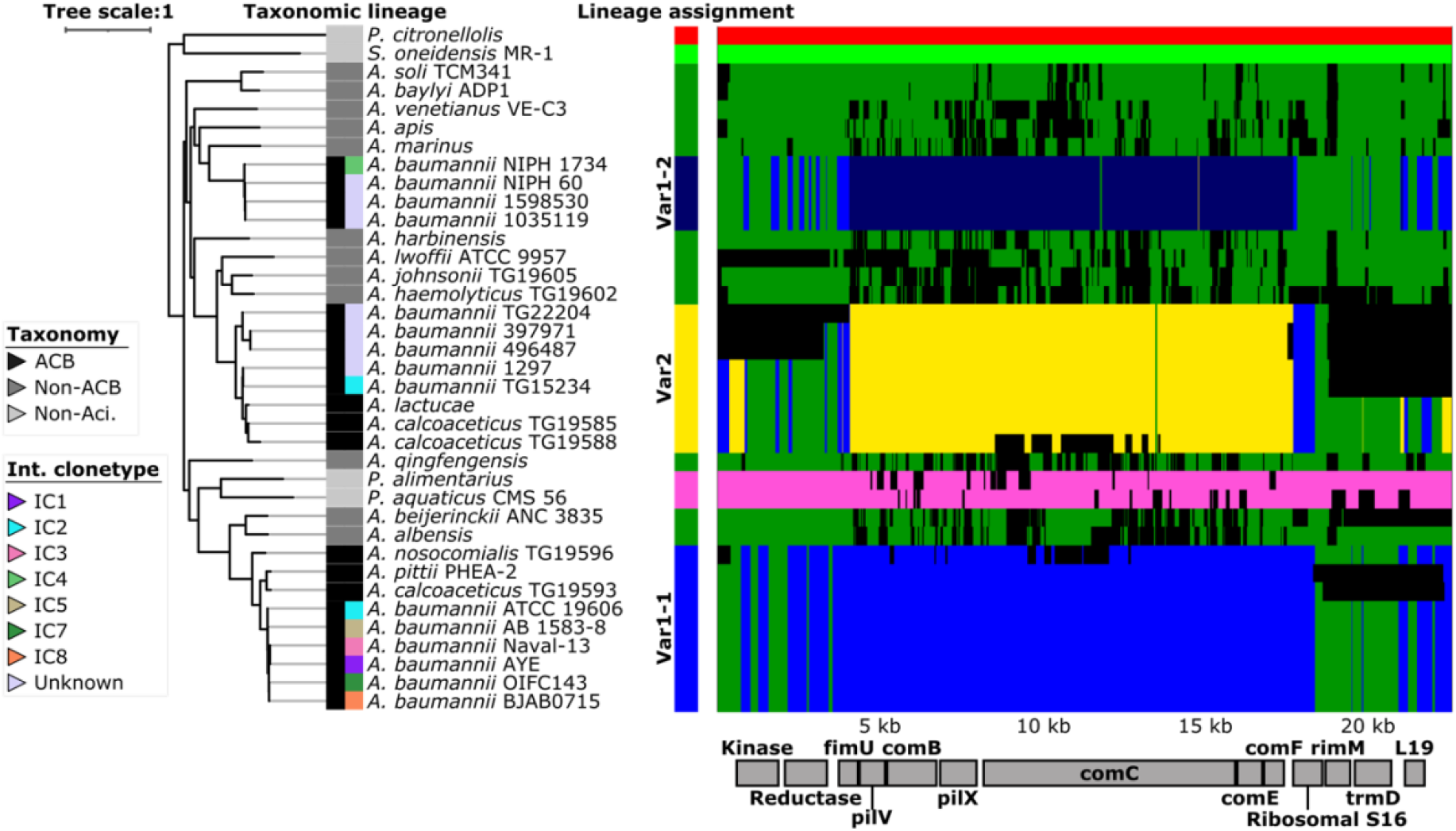
Patterns of ancestral recombination in the locus surrounding the *comC* gene. The main plot displays the inferred patterns of ancestral recombination for *Ab* ATCC 19606 ^T^ *comC* and the 12 flanking genes across 37 *Acinetobacter* isolates. Identity and order of the genes is indicated by the boxes below the plot. Each of the seven detected genetic lineages is represented by one color, and changes in color along the 13 genes indicate a recombination event. Alignment gaps or sites of unknown source of recombination are shown in black. The three genetic lineages represented in *A. baumannii* isolates are named Var1-1, Var1-2, and Var2 respectively (see main text). The maximum likelihood tree is based on the concatenated alignment of the seven T4aP proteins in this region. Branch length represent evolutionary time in substitutions per site. The taxonomic assignments together with the international clonetype (ICs) assignments for the *A. baumannii* isolates are indicated by the colored boxes left of the taxon labels. *P. citronellolis – Pseudomonas citronellolis; P. alimentarius* – *Psychrobacter alimentarius; P. aquaticus – Psychrobacter aquaticus; S. oneidensis – Shewanella oneidensis.*

Integrating the results of the evolutionary analyses with the domain architecture variant assignments for ComC and FimU provides substantial evidence that the entire cluster of T4aP associated genes represents an evolutionary cassette. The different domain architecture layouts of FimU and ComC characterize two main variants of this cassette, and an exchange of this cassette occurred at least twice during *A. baumannii* diversification.

### 3D modelling of ComC reveals variant-specific structural variation

In the highest resolving analysis, we assessed how the differences seen on the domain architecture level between ComC and FimU orthologs are reflected in the predicted 3D structures (Fig 4 and S9 Fig). ComC is characterized by the presence of two globular domains that are connected by a linker (Fig 4A). The C-terminal domain that harbors the Neisseria_PilC Pfam domain (PF05567; see Fig 1C) shows considerably little structural variation across the investigated proteins (see Fig 4B). In contrast, the N-terminal half is structurally less conserved. In *Ab* ATCC 19606 ^T^, this part of ComC is predicted to fold into an α/β doubly wound open twisted beta sheet conformation, which is surrounded by 7 parallel alpha helices arranged in a cylindrical conformation and an external alpha helix (Figure S10A). A similar layout is seen also in other ComC orthologs (S10 Fig B-C), and this fold agrees with previous structural characterization of the von Willebrand factor type A domain [50, 51], and of the VWA domain in integrin α II b (PDB: 3NIG). However, ComC_Var1_ is uniquely characterized by the presence of a finger-like protruding domain that carries a tyrosine (Tyr)-rich motif (Fig 4 and S10 Figs A-B). In ComC_Var1-1_, this finger is located between the alpha helices 3 and 4, whereas in ComC_Var1-2_ is located further towards the N-terminus between the first beta sheet and alpha helix 1 (Fig 4C and S10 Fig). Therefore, the two fingers are very likely of different evolutionary origins although their structural similarity and the shared presence of the Tyr-rich motif suggest that they originated from the same source (Fig. 4D-E). Neither ComC_Var2_ in *A. baumannii* or in *A. baylyi,* nor PilY1 in *P. aeruginosa* are in possession of a similar protrusion (Fig 4A-B). However, a structure comparison to proteins in PDB revealed a considerable diversity of ComC orthologs that is driven by lineage-specific insertions into the structural backbone that is formed by the VWA domain (S11 Fig).

**Figure 4.**
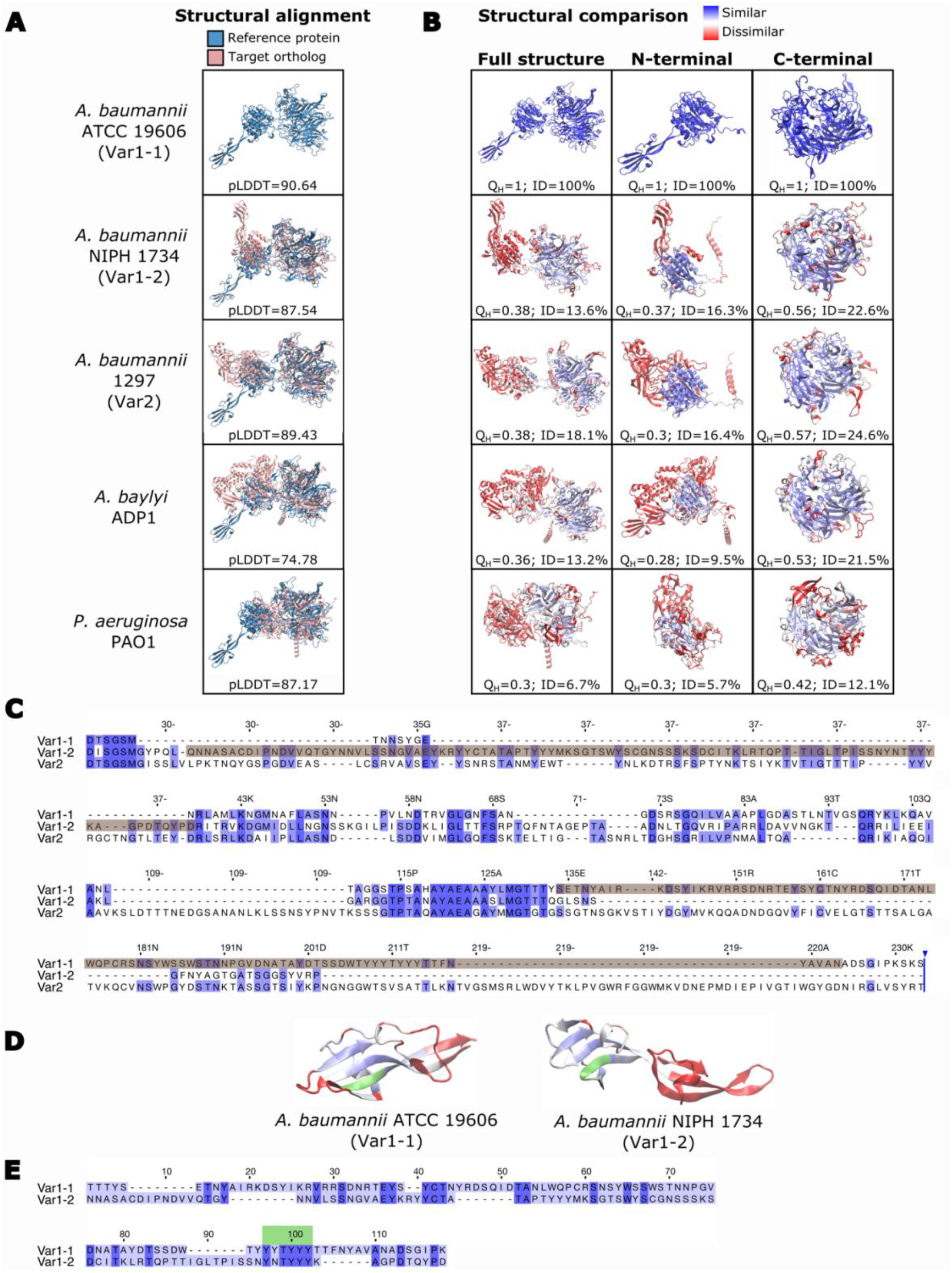
Variation of ComC in *A. baumannii.* (A) Alignments of the modelled 3D structures for *Ab* ATCC 19606 ^T^ ComC (blue) and the ComC of other bacterial isolates (red). The per-residue confidence for each modelled structure is given as the pLDDT value below the structures. Values >90 indicate a high accuracy and values between 70 and 90 a good accuracy of the prediction. (B) Structural similarity between ComC proteins shown in (A) and the reference protein in *Ab* ATCC 19606 ^T^. The structure of the target protein is colored with a gradient from blue (high conservation) to red (low conservation). Q_H_: pairwise structural conservation score ranging from 1 (structurally identical) to 0 (no similarity). ID: percent of sequence identity in the structural alignment. (C) Multiple sequence alignment of the N-terminal part of representatives for the three ComC variants in *A. baumannii.* The alignment covers the amino acids 24 – 233 of *Ab* ATCC 19606 ^T^ ComC. The sequences forming the finger-like protrusions in ComC_Var1-1_ and ComC_Var1-2_ are shaded in brown. The sequences for the three variants represent the corresponding species in shown in (A). (D) Structural similarity between the finger-like protrusions of ComC_Var1-1_ and ComC_Var1-2_. The color gradient from blue to red indicates decreasing similarity. The tyrosine-rich motif is highlighted in green. (E) Pair-wise sequence alignment between the sequences forming the finger-like protrusion in ComC_Var1-1_ and ComC_Var1-2_. Conserved residues are indicated in dark blue; the tyrosine-rich motif is shown in green.

### Functional role of ComC in *A. baumannii*

The *in-silico* analysis has provided substantial evidence for a hitherto unknown diversity of T4aP within *A. baumannii*. A prominent driver of this diversity is the pilus tip adhesin, ComC, and here specifically the N-terminal region that harbors a VWA_2 Pfam domain in ComC_Var1_. Eukaryotic proteins that incorporate VWA_2 domains participate in numerous biological functions among them cell adhesion and migration [52]. In line with this, we find a local structural similarity of *Ab* ATCC 19606 ^T^ ComC to mechanosensitive integrins, a superfamily of cell adhesion receptors in animals [53] (see S11 Fig). With the following experiments, we shed initial light on the functional relevance of the VWA_2 Pfam domain and of the finger-like structure therein. We created three different variants of ComC: the full-length version of *Ab* ATCC 19606^T^ *comC,* a truncated variant that lacks the VWA domain (*comC*Δ*VWA*), and a variant where we exclusively deleted the region that encodes the finger in ComC_Var1-1_ (*comC*Δ*166-256*). Note that a comparison of the predicted structures for ComC and for the ComCΔ166-256 mutant provided no evidence for a misfolding of the mutant (S12 Fig). The subsequent experiments were performed in a *comC* knock-out mutant of *Ab* AYE-T, because *Ab* ATCC 19606 ^T^ did neither twitch nor to take up environmental DNA in our hands.

We initially confirmed that *Ab* AYE-T Δ*comC* strain showed no noticeable piliation defect (Fig 5A). This finding is consistent with the observations that a *comC* deletion has no effect on piliation in *A. baylyi* [54] and in *Neisseria* [55], and we conclude that a deletion of *comC* does not interfere with piliation in any *A. baumannii* isolate. Subsequently, we investigated the role of ComC and of the VWA_2 Pfam domain in host cell adhesion (Fig 5B). Compared to wild-type *Ab* AYE-T, a *comC* knock out mutant (*Ab* AYE-T Δ*comC*) displayed a significantly reduced adhesion rate to HUVECs (n=4; t test: p<0.05). Complementing the mutant with the full length *comC* increased the adhesion rates significantly (n=4; t test: p<0.05), whereas no significant increase was observed when we used the either *comC*Δ*VWA* or *comCΔ166-256* for complementation. We next investigated the role of ComC in T4aP mediated twitching, and for natural transformation. *Ab* AYE-T Δ*comC* showed no twitching motility, and this phenotype was at least partly restored upon complementation with the full length *comC* (Fig 5C). Notably, neither of the truncated *comC* mutants could restore the capability to twitch to a noticeable extent. Similar to the effect on twitching motility, the deletion of *comC* abolished natural transformation (Fig 5D). Complementation with the full length *comC* almost fully restored the phenotype. Interestingly, this time the complementation with *comC*Δ*VWA* and *comC*Δ*166-256* also restored natural transformation, however with frequencies that are an order of magnitude below those of the full length *comC*.

**Figure 5.**
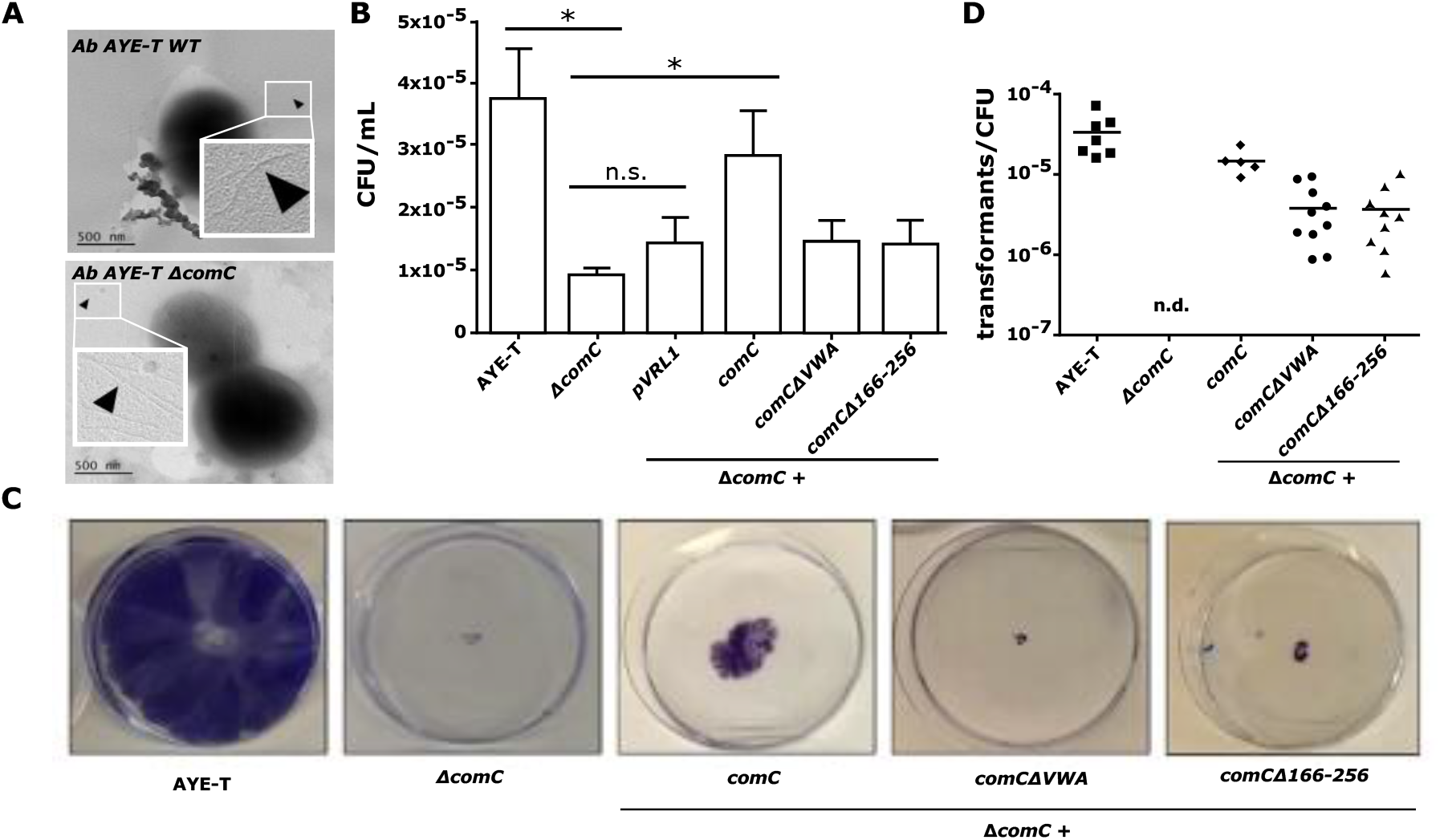
Functional characterization of ComC in *A. baumannii* AYE-T. (A) Representative electron micrograph of wild-type *A. baumannii* AYE-T cells (top) and of the Δ*comC* mutant (bottom). No piliation defect is seen for the mutant. A phenotyping of wild-type *A. baumannii* AYE-T, of the Δ*comC* mutant, and of the mutant complemented with the indicated constructs are shown in the following panels. (B) Adhesion to HUVEC cells. Bar height and error bar indicate the mean number and standard deviation of colony forming units (CFU) per mL (n=4; values are given as S2 Table). ‘*’ indicates a significant difference (one tailed t test: p<0.05). Complementation with the truncated *comC* mutants did not significantly increase adhesion rates. (C) Twitching motility. Cells were stained with 1% [w/v] crystal violet. (D) Natural transformation rates of the indicated *A. baumannii* cells was assessed using genomic DNA of rifampicin resistant *A. baumannii* ATCC 19606 ^T^. n.d. - not detected.

The experimental data provide first-time evidence that the N-terminal half of ComC and the VWA_2 Pfam domain therein play a critical role in T4aP mediated adhesion to HUVEC cells, for twitching and at least contribute to natural competence. While both twitching and natural competence was abolished upon *comC* deletion, the ability to adhere to HUVEC cells was only reduced. This is best explained by the effect of other adhesins, such as ATA [56], Csu fimbriae [57], or InvL [58], which all contribute to host cell adherence. Interestingly, we observe for all three tested functions the same effect when deleting only the finger-like protrusion that is characteristic for ComC_Var1-1_ instead of the entire VWA_2 domain. This provides initial evidence that at least part of the ComC_Var1-1_ function is conveyed by this finger, and that ComC variants lacking the finger differ also in function from ComC_Var1-1_.

## Discussion

Type IVa pili are prevalent in bacteria irrespective of their lifestyle where they convey a broad range of functions [24, 26]. In some and often only distantly related human pathogens, they represent key virulence factors [34, 35, 59, 60]. This indicates that the precise functions of T4aP have changed multiple times during evolution and probably as an adaptation to differing habitats and lifestyles. The genus *Acinetobacter* harbors environmental bacteria, bacteria that colonize various animals, as well as human pathogens [61]. This diversity in lifestyles provides an optimal setup for tracing genetic changes that underlie the functional diversification of T4aP that are used by the bacteria to interact with their environment.

The individual components necessary for building up the T4P machinery are almost ubiquitously present across the *Acinetobacter* diversity. This indicates that missing even one factor most likely renders the entire pilus dysfunctional. Along the same lines, it suggests that T4aP are essential for *Acinetobacter* fitness independent of both habitat and lifestyle. However, conspicuous differences between T4aP components of different *Acinetobacter* isolates emerged on the level of domain architectures. The connection between domain architecture of a protein and its function is well documented (e.g., [62–65]. Therefore, the differences for FimU and, more prominently for the pilus tip adhesin ComC, point towards a lineage-specific modification of T4aP function.

T4a pilus tip adhesins (T4a-PTA) have received considerable attention in various bacterial species, among them several human pathogens [55, 66–68]. Thus far, all investigated proteins share the presence of a Ca^2+^ binding domain in the C-terminal half (Pfam: Neisseria_PilC; PF05567), and assume similar roles in basal pilus function [55, 66, 69–72]. However, the precise functions of the N- and C-terminal globular domains differ among species. In *Neisseria,* the N-terminal half of PilC1 mediates host cell adherence [55, 66, 70, 71]. In *Legionella,* it is necessary for host cell invasion [73], and in *P. aeruginosa,* it has been associated with the mediation of surface adhesion, mechanosensing, and regulation of pilus retraction [38, 68]. In both *Legionella* and *P. aeruginosa,* the host cell adhesion function is mediated by the C-terminal half via an integrin-binding functionality [74].

Here, we have shown that the most prominent differences between the individual ComC orthologs both within *A. baumannii* and across the genus reside in the N-terminal part. Most *A. baumannii* isolates together with some members of the ACB clade feature a VWA_2 Pfam domain in this region. This domain was initially identified in the von Willebrand Factor of eukaryotes [75], and von Willebrand Factor A domains participate, among others, in cell adhesion and cell migration [76]. The presence of a VWA domain in a bacterial PTA was first described for the Pi-2a pilus in the Gram-positive *Streptococcus agalactiae,* a leading cause of sepsis and meningitis. As hypothesized from its function in eukaryotes, this domain indeed mediates host cell adhesion [77]. Subsequently, a VWA domain was also found in PilY1 of *P. aeruginosa* [38, 67, 78]. Therefore, our observation that VWA_2 Pfam domains are largely confined to ComC proteins of *Acinetobacter* pathogens in the ACB clade makes it tempting to postulate that the addition of a VWA domain to the PTA is at least one determinant that turns T4aP into a virulence factor. In this context, the recent finding is interesting that VWA domains can activate of macrophages, central regulators of airway inflammation [79, 80].

However, general features of a VWA domain are present also in ComC of a-pathogenic *Acinetobacter* isolates. This includes the characteristic metal ion-dependent adhesion (MIDAS) motif ((see S10 Fig; [81]) as well as extended stretches of structural similarity between the N-terminal region of ComC orthologs and the human von Willebrand Factor A (see S11 Fig). Why then were these domains not annotated with the VWA_2 Pfam domain? A sub-domain level analysis revealed differences between *Acinetobacter* ComC variants that likely result from lineage-specific insertions into a scaffold formed by a VWA domain (see S9 Fig and S10 Fig). This diversity is not reflected in the corresponding Pfam model, and thus domain instances with insertion patterns that are not represented in the training data of the Pfam model are likely to be missed. Although we found individual hits for these intervening sequences in the protein database (PDB), the similarity scores were low throughout, and thus the *in-silico* analyses currently provide no indication about the evolutionary origins and the likely functional roles of these insertions.

Testing the function of ComC_Var1-1_ *in-vivo* revealed that this protein is involved in host cell adhesion, twitching and DNA uptake, and that these functions are conveyed by the N-terminal half and of the VWA domain therein. Interestingly, a deletion of the finger-like protrusion in ComC_Var1-1_ was sufficient for impairing all three processes to an extent that is comparable to the deletion of the entire VWA domain. Because structural modelling revealed no indication that the deletion of the finger results in misfolding of ComC (see S12 Fig), these findings suggest a functional role of this structure. Still the evidences are only preliminary, and further analyses will be necessary to prove the involvement of this finger in ComC function. It will then also be interesting to see whether the Tyr-rich motif, that is present in the fingers of ComC_Var1-1_ and ComC_Var1-2_ has a functional role. Tyrosine assumes a broad spectrum of functions in natural systems, and short tyrosine rich peptides display a high propensity for self-assembly [82]. A functional role of this motif in ComC_Var1_ is therefore conceivable.

Next to ComC, also FimU displays a variation both in domain architecture and in the inferred 3D structure across the investigated isolates. The GspH Pfam domain, whose presence in FimUVar1 correlates well with the presence of ComC_Var1_, is exclusively found in bacterial proteins and it is characteristic of pseudopilins [83]. The absence of this domain in FimUVar2 is therefore intriguing, but in contrast to ComC its functional relevance is harder to predict. However, its tight association with ComC_Var2_ in one layout of the T4aP cassette hints towards a functional integration of the variants. It is thus conceivable that the exchange of this cassette, which happened at least twice during *A. baumannii* diversification created structurally, and most likely also functionally different T4aP that may have helped the bacterium to adapt to a specific environment.

The presence of structural variants of PilA (ComP in this study) has previously suggested a functional variation of T4aP in *Acinetobacter baumannii* [41]. Our data did not allow to reproduce this observation, because the structural variation among PilA proteins is not reflected in differences of their feature architectures (see Fig 1). However, reconciling the phylogenetic distribution of the PilA variants with those of the ComC and FimU variants reveals discrepant patterns. For example, while Ab ATCC 19606 ^T^ and Ab ACICU differ in their PilA structure [41], they both harbor the same *comC-fimU* cassette (see Fig 2). This strongly suggests that the evolutionary and functional plasticity of T4aP is substantially higher than anticipated. Thus, an essential part of how pathogenic *Acinetobacter* isolates interact with their environment in general, and with their host in particular, is largely uncharted. It will require highly resolved structural and functional studies to link the various T4aP layouts with lineage-specific differences in T4aP function and to assess the consequences for the bacterial phenotype.

## Conclusion

A broad range of bacterial taxa use Type IVa pili for the interaction with their environment, where their functional diversity has earned them the attribute “Swiss Army knife” among bacterial pili [34]. Here, we provided evidence that T4aP are substantially more diverse than it was hitherto appreciated. Already different isolates within *A. baumannii* seem to differ in their precise T4aP function, and thus in the way how they interact with the human host. This rapid change of pilus function seems to be achieved by an evolutionary concept that resembles an interchangeable tool system where the same handle can convey multiple functions depending on the precise layout of the tool head, here represented by the pilus tip adhesin ComC. Increasing the resolution to trace the functional modification of ComC on the subdomain level reveals the same concept. A conserved structural scaffold formed by the von Willebrand factor domain appears to be structurally and functionally modified by individual and lineage-specific insertions. On a broader scale, our findings suggest that a substantial extent of functional differences between bacteria isolates that is conveyed by changes on the domain- or sub-domain level rather than by the differential presence/absence of genes remains to be detected. Future comparative genomics approaches that aim to unravel the genetic specifics of pathogens should therefore best extend across different scales of resolution. The goal is to integrate lineage-specific differences on the level of gene clusters, genes, domain- and sub-domain architectures into a comprehensive reconstruction of molecular evolution in a functional context.

## Materials and Methods

### Data collection

Genome assemblies for 855 isolates from the genus *Acinetobacter* were retrieved from the RefSeq database release 204 [84]. 27 isolates of closely-related γ-proteobacteria and two representatives from the genus *Neisseria* were added as outgroups. The full taxon list is provided as S3 Table.

### Annotation of protein domains

Pfam and SMART domains were annotated with *hmmscan* from the HMMER package [85] using Pfam release 32.0 [86] and SMART release 9 [87]; we further annotated signal peptides with SignalP [88], transmembrane domains with tmhmm [89], coiled-coils with COILS2 [90], amino acid compositional bias with fLPS [91] and low complexity regions with SEG [92].

### Domain-architecture-aware ortholog search

Targeted ortholog searches were performed with fDOG v0.0.8 (www.github.com/BIONE/fDOG) [93] using *A. baumannii* ATCC 19606 ^T^ as the reference taxon. Domain architectures similarities between orthologs were computed with the software package FAS v1.3.2 (https://github.com/BIONF/FAS) [65] that is integrated into the fDOG package. Phylogenetic profiles were visualized and analyzed using PhyloProfile v1.8.0 [94]. The taxa represented in the phylogenetic profiles were ordered with increasing phylogenetic distances from *A. baumannii* ATCC 19606 ^T^ using the *Acinetobacter* phylogeny in [20] and the γ-proteobacteria tree from [95].

### Phylogenetic analyses

Protein sequences were aligned using MAFFT v7.394 with the linsi method [96]. Phylogenetic gene trees were computed using RAxML v8.2.11 [97] with the rapid bootstrapping algorithm and 100 replicates. The WAG model [98]was selected via the PROTGAMMAAUTO option of RAxML as the best-fitting substitution model. Trees were visualized and annotated using the iTOL website [99]. Alternative topology testing was performed using the AU test [49] as implemented into the software MATT (https://github.com/BIONF/MATT).

### Detection of ancestral recombination

A subset of 33 *Acinetobacter* isolates together with four outgroup taxa where selected to represent both the phylogenetic diversity and the different ComC layouts in our data. For each taxon, we extracted the genomic region between 5,000 bp upstream and 3,500 bp downstream of *comC* and aligned the sequences with MAFFT v7.394. The detection of individual genetic lineages together with the prediction of ancestral recombination events was done with fastGEAR [100] using default parameters. Shared synteny analyses of the genes annotated in the region across the investigated taxa was assessed using the software Vicinator (https://github.com/ba1/Vicinator) with the orthology assignment from fDOG [20].

### 3D structure analysis

AlphaFold2 [101] together with the reduced PDB database was used to model the 3D structures for the following proteins: ComC – *Ab* ATCC 19606 ^T^ (WP_085940514.1), *Ab* 1297 (WP_024436449.1), *Ab* NIPH 1734 (WP_004745407.1), *A. baylyi* ADP1 (WP_004923840.1), *A. qingfengensis* (WP_070071002.1); PilY1 – *Pseudomonas aeruginosa* PAO1 (NP_253244.1) PilY – *Legionella pneumophila* (WP 229293926.1); PilC –*Neisseria gonorrhoeae*. The structures for *Legionella. pneumophila* PilY and *Neisseria gonorrhoeae* PilC were retrieved from existing predictions in UniProt (accessions Q5ZXV3 and Q5FAG7, respectively). Models were visualized in VMD [102], and structural conservation between ComC orthologs and *Ab* ATCC 19606 ^T^ ComC were analysed with the MultiSeq extension from VMD [103] calling the structural alignment tool STAMP [104]. Subdomains and structural similarities to other proteins in the PDB database were identified using the Vector Alignment Search Tool: VAST [105].

### Culture conditions of bacterial strains and cell lines

All *A. baumannii* strains used for experiments in this study are listed in Table 2. Strains were grown in Luria-Bertani medium (LB) at 37 °C with 50 μg ml^-1^ kanamycin or 100 μg ml^-1^ gentamicin when needed. Human umbilical vein endothelial cells (HUVECs) were extracted from fresh cord veins and cultivated in endothelial growth medium. Medium was supplemented with growth factor mix and 10% fetal calf serum.

**Table 2.**
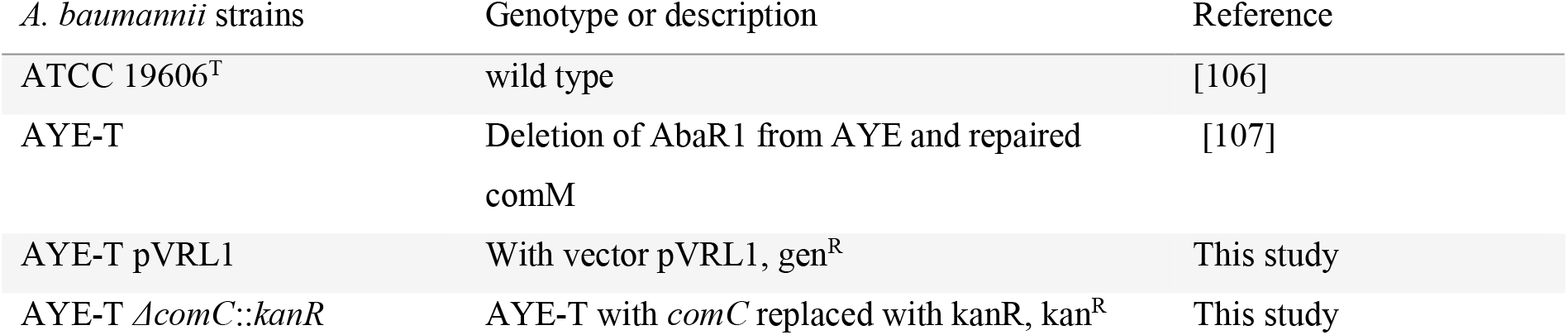
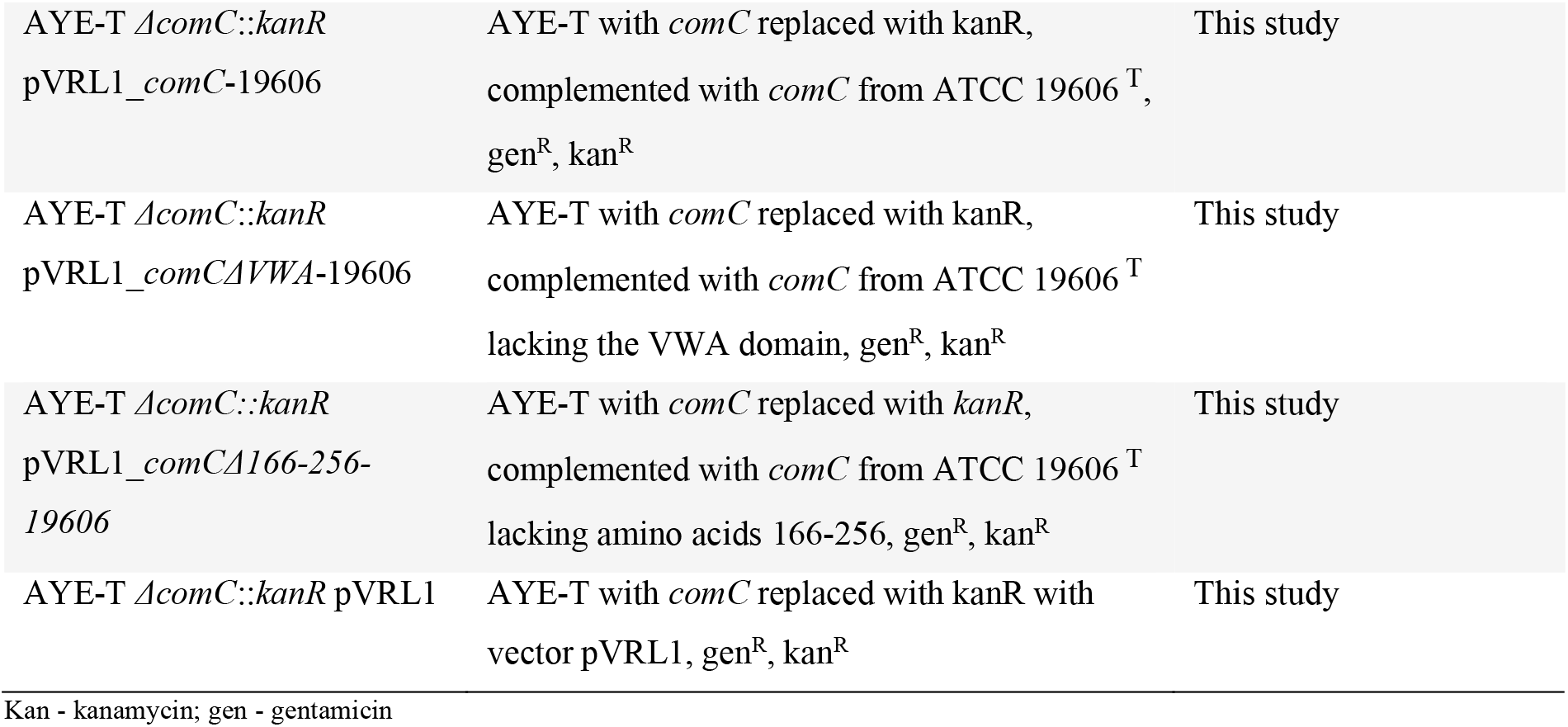
Bacterial strains used in this study.

### Generation of mutants and complementation analysis

The *ΔcomC::kanR A. baumannii* AYE-T deletion mutant was generated as described by Tucker, Nowicki (108) using primers 1-10 (S4 Table). All mutants were verified by sequencing. Three different constructs were used to complement *Ab* AYE-T *ΔcomC::kanR:* (i) full length ComC – The *comC* gene plus 700 bp of the upstream region were amplified from chromosomal DNA of *A. baumannii* ATCC 19606^T^ using primers 11-12 (S4 Table). The amplicon was integrated into pVRL1 [109] using KpnI and XhoI resulting in plasmid pVRL1_*comC-*19606. (ii) ComC lacking the VWA-domain: *pVRL1_comCΔVWA-*19606 was amplified from pVRL1_*comC*-19606 using primers 13-14 followed by blunt end ligation. (iii) ComC lacking the finger-like protrusion: amino acids 166-256 were deleted using plasmid pVRL1_*comC*-19606 and primers 15-16 resulting in plasmid pVRL1_*comCΔ166-256*-19606. Plasmids were transferred into the mutant *via* electroporation.

### Natural transformation

Wild type and mutant strains were grown in LB medium overnight at 37 °C and diluted to OD600 0.01 with phosphate buffered saline (PBS). Equal amounts of the bacterial suspension and DNA (100 ng/μl genomic DNA of rifampicin resistant *A. baumannii* ATCC 19606 ^T^) were mixed and 2.5 μl of the mixture were applied onto 1 ml of freshly prepared transformation medium (5 g/L tryptone, 2.5 g/L NaCl, 2 % [w/v] agarose) in 2 ml reaction tubes. After incubation for 18 hours at 37 °C, cells were resuspended from the medium with PBS. Transformants were selected by plating on selective agar (rifampicin 20 μg ml^-1^).

### Twitching motility

Twitching medium (5 g/L tryptone, 2.5 g/L NaCl, 0.5 % agarose) was inoculated by stabbing one bacterial colony through the agar to the bottom of the petri dish. Plates were sealed with parafilm to prevent desiccation and incubated at 37 °C for three days. To visualize the cells at the bottom of the petri dish, the agar layer was removed, and cells were stained with 1% [w/v] crystal violet.

### Piliation analyses by electron microscopy

To analyze the piliation phenotype, cells were grown on LB agar plates at 37 °C overnight. Cells were prepared for electron microscopy and visualized as previously described [110]. Shadowing of the cells was carried out in an angle of 20° (unidirectional) and with a thickness of 2 nm platinum/carbon.

### Analysis of bacterial adhesion to human endothelial cells

Primary human umbilical cord vein cells (HUVECs) were cultivated in endothelial growth medium supplemented with growth factor mix (ECGM, Promocell) and 10% fetal calf serum (FCS, Sigma-Aldrich) in collagenized 75 cm^2^ cell culture flasks using a humidified incubator with a 5% CO2 atmosphere at 37 °C. HUVECs were seeded into six-well plates and infected with *A. baumannii* (MOI 50) for three hours. The supernatant was removed and cells with adherent bacteria were washed with PBS and detached using a cell scraper. Thereafter, adherent bacteria were quantified by plating serial dilution series. Visualisation and quantification of bacterial adhesion to human endothelial cells was done by fluorescence microscopy as described in [56].

## Declarations

### Data availability statement

The authors confirm that all data underlying the findings are fully available without restriction. All numerical data that underlies figures and/or summary statistics is within the manuscript and its Supporting Information files. Genomic data analyzed in this study are publicly available in the NCBI Reference Sequence Database (RefSeq). For each selected genome, the corresponding assembly accession is listed in S3 Table. We obtained protein sequences and coding sequences (*.faa and *cds_from_genomic.fna), and genome sequence (*_genomic.fna) files from the database [ftp://ftp.ncbi.nlm.nih.gov/refseq/release/release-catalog/archive/]. The domain architecture-aware phylogenetic profiles for the *Ab* ATCC 19606 ^T^ T4aP components are available via figshare: https://figshare.com/articles/dataset/_/21964535. A high-resolution version of the tree shown as S6 figure is available via figshare: https://figshare.com/articles/figure/_/21967694.

### Competing interests

The authors declare that they have no competing interests.

### Funding

This study was supported by a grant by the German Research Foundation (DFG) in the scope of the Research Group FOR2251 “Adaptation and persistence of A. baumannii.” Grant ID EB-947 285-2/2 to IE, AV 9/7-2 to BA, GO 2491/1-2 to SG. BA additionally acknowledges financial support by the Deutsche Forschungsgemeinschaft via AV 916-2, and IE was further supported by the Research Funding Program Landes-Offensive zur Entwicklung Wissenschaftlich-ökonomischer Exzellenz (LOEWE) of the State of Hessen, Research Center for Translational Biodiversity Genomics (TBG).

## Acknowledgments

The authors wish to thank all researchers for making annotated genome sequences available to the public domain. We thank M. Linder from the Max-Planck-Institute of Biophysics, Frankfurt am Main, for the support of electron microscopical analyses, X. Charpentier from the Claude Bernard University, Lyon for providing strain *A. baumanii* AYE-T, Holger Bergman and Sachli Zafari for their contribution in the early phase of the project, and Felix Langschied for helpful discussion and critically reading of the manuscript.

## Author Contributions

Conceptualization: Ingo Ebersberger

Data curation: Ruben Iruegas

Formal analysis: Ruben Iruegas, Ingo Ebersberger

Funding acquisition: Beate Averhoff, Stephan Göttig, Ingo Ebersberger

Investigation: Ruben Iruegas, Katharina Pfefferle, Stephan Göttig, Ingo Ebersberger

Methodology: Ruben Iruegas, Beate Averhoff, Ingo Ebersberger, Stephan Göttig

Project administration: Ingo Ebersberger.

Resources: Beate Averhoff, Stephan Göttig, Ingo Ebersberger

Supervision: Beate Averhoff, Ingo Ebersberger

Validation: Ingo Ebersberger.

Visualization: Ruben Iruegas, Ingo Ebersberger.

Writing – original draft: Ruben Iruegas, Ingo Ebersberger.

Writing – review & editing: Ruben Iruegas, Katharina Pfefferle, Stephan Göttig, Beate

Averhoff, Ingo Ebersberger

## Supplementary information

### Supplementary Figures

**S1 Figure. Blastp hits of *A. baumannii* ATCC 19606^T^ ComC (WP_085940514.1) in the NCBI nrProt database**

**S2 Figure. Domain architecture comparison of Ab ATCC 19606 T4aP components whose differences between orthologs do not include Pfam or SMART domains.** For the three represented components (A-C), the differences in domain architectures between the protein in Ab ATCC 19606 and their orthologs in other isolates are limited to coiled coil regions, low complexity regions (LCR) and regions with a compositional bias (FLPS) (see main text, Fig. 1). Reference proteins shown in the first column are compared against their orthologs in *Acinetobacter soli* CIP 110264 (GCF_000368705.1). TMM – Transmembrane domain; Gln – Glutamine; Ile – Isoleucine.

**S3 Figure. E-value distribution PilX_N Pfam domain annotations in PilX orthologs.** Only a subset of PilX orthologs was annotated with a PilX_N Pfam domain (PF13681) whereas the domain architecture of the remaining orthologs comprise only a transmembrane domain (TMM) and a low complexity region (LCR) (A). The box plot shows the e-value distribution for the annotated PilX_N Pfam domains. Most values are high and close to the inclusion limit of 0.001. Thus, the variation in domain architectures between PilX orthologs is likely due to chance effects affecting the accurate annotation of the PilX Pfam domain.

**S4 Figure. Characterization of the pilin FimT**. (A) Pair-wise domain-architecture similarities between *Ab* ATCC 19606^T^ FimT and its 864 orthologs across the analyzed taxa [65]. The upper block indicates the similarity values penalizing the absence of an Ab ATCC 19606 domain in the ortholog (FAS_F). The consistent coloring in blue indicates that the overall domain architecture similarity is high. The lower block penalizes the absence of a domain that is seen in the ortholog but that is absent in *Ab* ATCC 19606^T^ FimT. Most orthologs are colored in pink, which indicates that they possess a domain that is not seen in *Ab* ATCC 19606 ^T^ FimT. (B) Pairwise domain architecture comparison between FimT in *Ab* ATCC 19606 ^T^ (WP_000477156.1) and its ortholog in *Ab* ACICU (WP_000477147.1). This reveals the absence of an N_methyl Pfam domain (purple) in the N-terminal region of *Ab* ATCC 19606 ^T^ FimT. (C) Pairwise sequence alignment between FimT in *Ab* ATCC 19606 ^T^ and in *Ab* ACICU. Conserved residues are highlighted in dark blue; substitutions are shown in light blue. FimT of *Ab* ATCC 19606 ^T^ carries a threonine at position 9 in the alignment. (D) The HMM weblogo of the N_methyl domain reveals that position 9 is typically occupied by either leucine or isoleucine, which explains why no N_methyl domain was annotated in FimT of *Ab* ATCC 19606 ^T^. ACB – *Acinetobacter calcoaceticus-baumannii* complex; γ – *γ-proteobacteria;* β – *β-proteobacteria.*

**S5 Figure. Domain architecture changes of ComC and FimU within the genus Acinetobacter.** (A) The two main domain architecture (DA) variants of ComC and FimU, respectively, in the genus *Acinetobacter*. (B) The DA represented in *Ab* ATCC 19606^T^ is dominant in the ACB complex. “FAS” displays the distribution of domain architecture similarity scores between ComC in Ab ATCC 19606 and its orthologs separately for isolates inside (red) and outside (blue) of the ACB complex. FAS scores range from a maximum of 1 (identical architectures) to 0 (no shared domains) [1]. The protein of *Ab* ATCC 19606 is used as reference. The asterisk marks the FAS score mean. The average number of instances per proteins (IPP) for the individual domains is represented by the bar plot. Left plot: ComC; right plot: FimU. Pfam domains: VWA_2 - PF13519; Neisseria_Pil_C - PF05567; N_Methyl - PF07963; GspH - PF12019.

**S6 Figure. Maximum likelihood phylogeny of ComC.** Annotations on the right show the assignments of taxonomic groups, protein variants of ComC and FimU, and the cassette type. A high-resolution version of this tree is available via figshare: https://figshare.com/articles/figure/_/21967694

**S7 Figure. Domain architecture diversity for ComC and FimU in the genus *Acinetobacter.*** The tree shows a selection of taxa from the full set that represents the diversity of domain architectures of ComC and FimU. The color code of the leaf labels resembles that of Fig. 3 in the main text. The domain architectures of ComC and FimU are given next to the taxon names.

**S8 Figure. Gene order conservation analysis in the genomic region harboring ComC and FimU.** Each line in the plot represents the bacterial isolate that is indicated to the left. The color coding of the taxon labels corresponds to their lineage assignment in Fig. 3 of the main text. Each box represents an ortholog to one of the 13 genes in the ComC/FimU region of *Ab* ATCC 19606 (see main Fig. 3) and the gene identity is given by the box label. The order of boxes corresponds to the order of genes in the genome of the respective isolate. *comC* orthologs are marked in white. Green and red boxes identify genes that are upstream and downstream of *comC* in *Ab* ATCC 19606, respectively. The number of genes separating any two of the boxed genes or that are placed upstream or downstream up to the scaffold end (indicated by yellow bars) are given in parenthesis. The plot confirms that the gene order in this region is conserved throughout the genus with very few exceptions.

**S9 Figure. Comparison of two FimU variants.** (A) Comparison of the modelled 3D structures of FimUVar1 (*A. baumannii* ATCC 19606^T^) and FimUVar2 (*A. baumannii* 1297). The structures are colored according to their mutual structural similarity from high (blue) to low (red). Red colored alpha helices in both structures correspond to the signal peptide. (B) Pairwise sequence alignment of the two FimU variants. Residue numbering refers to FimU in *Ab* ATCC 19606.

**S10 Figure. Structural fold of the N-terminal domain of the ComC orthologs.** Alpha helices (orange) and beta strands (blue) are shown in the 3D cartoon representation (left) and in the secondary structure topology plots of sample proteins (right). Highlighted are the MIDAS motif (yellow), tyrosine-rich motif (light green), subdomains (brown and bright blue), and external folds (grey, purple, dark green). *A. baumannii* ATCC 19606 (A) and *A. baumannii* NIPH 1734 (B) feature an antiparallel beta sheet finger-like protrusion (brown). (C) The structure of *A. baylyi* contains additional folds (dark blue, purple, green, and brown) that are highly disordered and form a planar shield that surrounds the MIDAS motif.

**S11 Figure. VAST search for regions of structural similarity in the N-terminal half of ComC for selected bacterial isolates.** For each of the analyzed proteins, the following information is given: Top left – linear domain-representation of the ComC N-terminal region, where different domains are indicated by different colors. The location of the domain in the 3D structure model to the right is given by the correspondingly colored region. Bottom - Segments sharing a significant similarity to an existing structure in the PDB database are identified by arrows beneath the linear domain representation, and we limited the information to the three highest scoring hits. Information about the hit protein is given to the right. (A) *A. baumannii* ATCC 19606 (ComC_Var1-1_); (B) *A. baumannii* NIPH 1734 (ComC_Var1-2_); (C) *A. baumannii* 1297 (ComC_Var2_); (D) *A. baylyi* ADP1; (E) *A. qingfengensis;* (F) *Pseudomonas aeruginosa;* (G) *Legionella pneumophila;* (H) *Neisseria gonorrhoeae.* The analyses reveal that ComC orthologs differ by the presence of sequences with weak structural similarities to a diverse set of proteins in PDB. This indicates that a structural backbone of a VWA domain is modified, in a lineage-specific way, by insertions of different origins.

**S12 Figure. Comparison of the predicted structures of Ab ATCC 19606 ComC and of Ab ATCC 19606 ComC □ 156-266.** The predicted structures of Ab ATCC 19606 ComC and of the mutant lacking the finger like protrusion are shown in panels (A) and (B), respectively. The hatched box in (A) indicates the finger in the wild-type protein. A structural alignment of the two N-terminal globular domains is shown in (C), where structurally conserved regions are shown in blue.

### Supplementary Tables

**S1 Table. Pfam domains annotated in type IVa pilus proteins of *A. baumannii* ATCC 19606.**

**S2 Table. Results of the adhesion experiments.** Values are given in colony forming units per mL

**S3 Table. List of taxa analyzed in this study.** The list is provided in the file S3_table.xlsx (.xlsx)

**S4 Table. Primers used in this study.**

**Supplementary Data**

**S1 Data. Orthology-based phylogenetic profiles for the *A. baumannii* ATCC 19606 T4aP components.**

## Notes

### Competing Interest Statement

The authors have declared no competing interest.

